# Engraftment of allotransplantated tumour cells in adult *rag2* mutant *Xenopus tropicalis*

**DOI:** 10.1101/2021.11.15.468684

**Authors:** Dieter Tulkens, Dionysia Dimitrakopoulou, Tom Van Nieuwenhuysen, Marthe Boelens, Suzan Demuynck, Wendy Toussaint, David Creytens, Pieter Van Vlierberghe, Kris Vleminckx

**Affiliations:** Department of Biomedical Molecular Biology, Ghent University, Ghent, Belgium; Cancer Research Institute Ghent (CRIG), Ghent, Belgium; Laboratory of Myeloid Cell Ontogeny and Functional Specialization, VIB-UGent Center for inflammation Research, Ghent, Belgium; Department of Pathology, Ghent University and Ghent University Hospital, Ghent, Belgium; Department of Biomolecular Medicine, Ghent University, Ghent, Belgium

## Abstract

Modelling human genetic diseases and cancer in lab animals has been greatly aided by the emergence of genetic engineering tools such as TALENs and CRISPR/Cas9. We have previously demonstrated the ease with which genetically engineered *Xenopus* models (GEXM) can be generated. This included the induction of autochthonous tumour formation by injection of early embryos with Cas9 recombinant protein loaded with sgRNAs targeting multiple tumour suppressor genes. What has been lacking so far is the possibility to propagate the induced cancers via transplantation. In this paper we describe the generation of a *rag2*^*-/-*^ knock-out line in *Xenopus tropicalis* that is deficient in functional T- and B-cells. This line was validated by means of an allografting experiment with a primary *tp53*^*-/-*^ donor tumour. In addition, we optimized available protocols for sub-lethal gamma irradiation of *X. tropicalis* froglets. Irradiated animals also allowed stable, albeit transient, engraftment of transplanted *tp53*^*-/-*^ tumour cells. The novel *X. tropicalis rag2*^*-/-*^ line and the irradiated wild type froglets will further expand the experimental toolbox in this diploid amphibian, and help to establish it as a versatile and relevant model for exploring human cancer.

## Introduction

The earliest transplantation of human primary tumour cells in mammalian hosts was described by Dr. Harry S. N. Greene (1938). Gradually during the last decades, tumour transplantation has been recognized as an indispensable tool in the cancer research field and has been successfully performed not only in mammalian species such as mice [reviewed by Sharkey & Fogh (1984)] but also in non-mammalian vertebrates like zebrafish [reviewed by Gansner et al. (2017)]. Cancer immunoediting, and more specifically cancer immunosurveillance, is an important process that can severely hamper engraftment of tumours in immunocompetent hosts (Dunn et al., 2002). In order to escape from this phenomenon either inbred or immunodeficient animals are required, thus allowing stable tumour progression after transplantation. Researchers working with mice were able to generate, amongst others, the ‘nude mice’ (lacking the thymus and thus functional T-cells), the NOD-SCID and SCID-beige mice that are deficient in both the T- and B-cell pool, and finally the NSG or NOG mice that additionally lack functional NK cells (Yoshida, 2020). More recently zebrafish have joined the field. Several protocols and resources are available in this species to achieve stable engraftment of transplanted cells such as for example sub-lethal irradiation (Traver et al., 2004), the use of a *rag2*^*E450fs*^ immunocompromised animals (Tang et al., 2014) and the use of syngeneic zebrafish lines, *e*.*g*. the CG1-strain (Smith et al., 2010). Furthermore, for xenograft experiments this species holds great promise as the transparent *casper* strain allowed the tracing and functional characterization of fluorescently labelled human tumour cells (White et al., 2008). Most recently the Langenau lab generated adult *prkdc*^-/-^, *il2rgα*^-/-^ immunocompromised zebrafish in the casper-strain that allowed robust engraftment of human cancer cells (Yan et al., 2020).

*Xenopus*, like zebrafish enjoys transparency in embryonic stages, allowing tracing of fluorescently labelled cells. Besides, the *Xenopus* innate and adaptive immune cells and mechanisms show high conservation with their respective mammalian counterparts (Banach & Robert, 2017). Despite the emergence of *Xenopus tropicalis* as a cancer model, thanks to the ease with which genetically engineered *Xenopus* models (GEXM) can be generated, so far experiments with tumour transplantations have not been documented for this species. Transplantations of *X. laevis ff-2* lymphoid tumour cells in inbred MHC homozygous *ff X. laevis* animals have led to the interesting finding that grafts are accepted in transplanted tadpoles but rejection is present in transplanted adults (Robert et al., 1995, 1997). This phenomenon is believed to be due to the second histogenesis present in the thymus during and after metamorphosis (Robert et al., 1995, 1997). Recently, Rollins-Smith & Robert (2019) described a protocol to induce lymphocyte deficiency by subjecting *X. laevis* frogs to sub-lethal gamma irradiation. Another study (Rau et al., 2001) showed engraftment successes after transplanting the 15/0 lymphoid tumour line (from a spontaneous *X. laevis* thymoma) in *X. laevis* irradiated hosts. We describe here the generation and validation of a novel immunodeficient *rag2*^*-/*-^ *X. tropicalis* line, suitable for transplantation experiments. Furthermore, we optimized and validated current available protocols for transplanting primary *Xenopus* tumours, for the first time, in irradiated *X. tropicalis* hosts. We believe these robust tools will be of high value for *Xenopus* tumour transplantation experiments and tumour immunity studies in general.

## Results

### Generation of *rag2*^*-/-*^ line

In order to generate a *X. tropicalis rag2*^*-/-*^ line, an sgRNA was designed targeting the first fifth of the *rag2* single exon gene. Wild type embryos were injected with a mixture of the selected sgRNA and Cas9 recombinant protein (Fig. 1A). To analyse editing efficiency, stage NF 41 embryos were lysed and genotyped. Amplicon deep sequencing (MiSeq™ System – Illumina) of the targeted region in the *rag2* gene revealed a major inclusion of a 4 bp deletion, which is in correspondence with what is predicted by the inDelphi CRISPR repair outcome prediction algorithm (Shen et al., 2018). Correlation analysis revealed a significant high overall correlation between predicted and endogenously observed frequencies of variant calls (Pearson r = 0.9886, *p* < 0.0001) (Fig. 1B) confirming previous findings proposing inDelphi as suitable method for predicting CRISPR/Cas9 induced repair outcomes in *X. tropicalis* (Naert, Tulkens, et al., 2020). For obtaining homozygotes (see schematic Fig. 1A), first, crispant mosaic mutant animals were raised until adulthood, outcrossed with wild type animals and checked for germline transmission in the progeny. Heterozygote *rag2*^*+/mut*^ animals were subsequently intercrossed and homozygote *rag2*^*mut/mut*^ animals were selected using a mixed Heteroduplex Mobility Assay (mHMA) genotyping technique (Foster et al., 2019) (Fig. 1C top). Sanger sequencing confirmed biallelic presence of a 4 bp deletion in homozygous mutant animals (Fig. 1C bottom). This deletion induces a frameshift after amino acid 91 resulting in a non-functional protein. Therefore, these animals are further referred to as *rag2*^*-/-*^.

**Figure 1.**
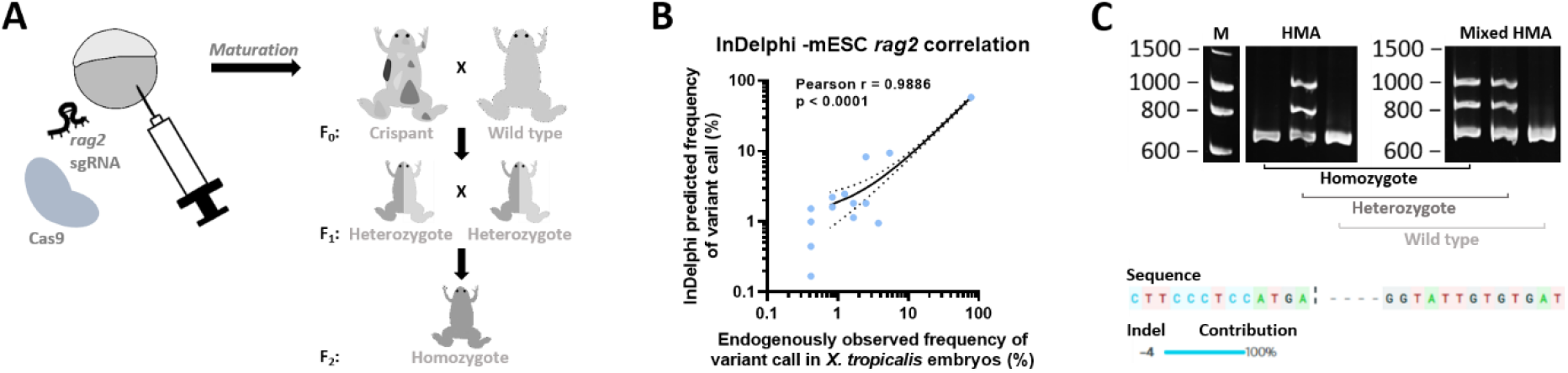
Generation of the *X. tropicalis rag2*^*-/-*^ knock-out line. **(A)** Embryos were injected with an sgRNA targeting the *rag2* gene along with Cas9 protein. When sexually mature, animals were outcrossed to wild types to obtain heterozygous animals that were subsequently incrossed to obtain *rag2* homozygous mutant animals in the F_2_ generation. **(B)** Scatter plot showing correlation between *in vivo* observed mutational CRISPR repair outcomes in injected embryos (x-axis) versus predicted outcomes using the inDelphi algorithm tool (y-axis). Dashed lines show the 95% confidence interval corresponding to the best-fit linear regression line (solid line). **(C)** Images taken from DNA electrophoresis gels after performing a normal HMA (left) and mixed HMA (right). Normal HMA included heating of the unknown PCR amplicons followed by slowly cooling and loading on the gel, while for mixed HMA, unknown PCR samples were first mixed with wild type *rag2* amplicons after which the normal HMA was performed. Multiple bands present in both gels indicate heterozygous animals, while extra bands appearing after performing the mixed HMA (right gel) relate to homozygous mutant animals. Absence of any extra bands is indicative of wild type animals.

### Transplantation of *X. tropicalis tp53*^*-/-*^ tumour in an *X. tropicalis rag2*^*-/-*^ adult

To assess transplantation potential in the novel *rag2*^*-/-*^ line, a thymic tumour originating from an adult *tp53*^*-/-*^ animal from a previous study (Naert, Dimitrakopoulou, et al., 2020) was isolated (Fig. 2A). Two parallel transplantations were performed: 5×10^6^ tumour single cells were transplanted intraperitoneally (IP) in a *rag2*^*-/-*^ and a wild type adult as illustrated in Fig. 2B. Ten weeks post transplantation the *rag2*^*-/-*^ transplanted animal showed obvious signs of lethargy, while the transplanted wild type showed no signs of discomfort. A clear externally visible outgrowth was present in the *rag2*^*-/-*^ animal close to the transplantation injection site (Fig. 2C). Upon dissection multiple sites of engraftment were observed on the abdominal muscle wall and in the peritoneal cavity (Fig. 2D). Histopathological analysis of the tumours revealed presence of both epithelial and mesenchymal cell clusters, thereby showing morphological similarities to the donor tumour (Fig. 2E, top). Interestingly, multiple zones with neovascularization were present in these tumour engraftment sites (Fig. 2E, top). In addition, immunohistochemistry showed high proliferative capacity in both donor and engrafted tumours as indicated by PCNA immunostaining (Fig. 2E, bottom). Finally, the in-house developed mixed HMA method confirmed the inclusion of the same *tp53* mutational variant, present in both the donor and the engrafted tumour (Fig. 2F). These data show that adult *rag2*^*-/-*^ knock-out *X. tropicalis* allows stable allografting of transplanted GEXM-derived tumour cells.

**Figure 2.**
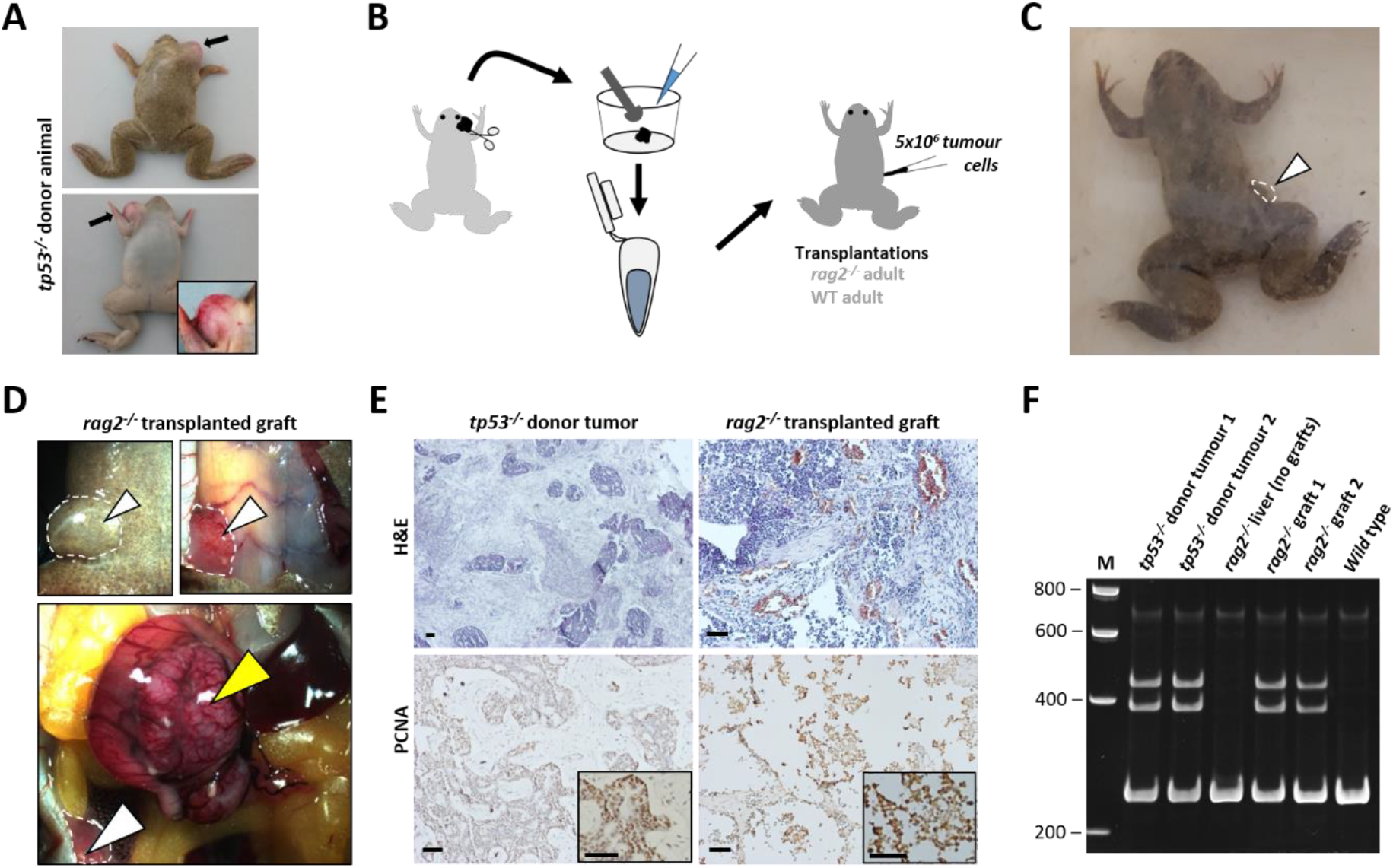
Validation of allografting in *X. tropicalis rag2*^*-/-*^ animals. **(A)** *tp53*^*-/-*^ donor animal harbouring a thymic tumour (black arrows). **(B)** Transplantation strategy including single cell generation using a 40 μm strainer followed by IP injections in a *rag2*^*-/-*^ adult and a wild type adult control (both 5×10^6^ live cells). **(C)** A *rag2*^*-/-*^ transplanted animal with visual subcutaneous outgrowth close to the injection site (white arrow, white dashed line) 10 weeks post-transplantation. **(D)** Dissection microscopy images (ventral view) of *rag2*^*-/-*^ transplanted animal showing external (top left) and internal (top right & bottom) views of the engrafted tumour at the injection site (white arrowheads, white dashed line) with an additional tumour mass in the intestinal mesenterium (yellow arrowhead). **(E)** H&E and IHC stained sections from primary tumour in *tp53*^*-/-*^ donor animal and the tumour graft in the *rag2*^*-/-*^ animal transplanted with the *tp53*^*-/-*^ tumour cells. **(F)** DNA electrophoresis gel image after performing a mixed HMA (for *tp53* gene) on DNA from two *tp53*^*-/-*^ tumour samples (donor animal), liver (without grafts) and two tumour grafts obtained from the transplanted *rag2*^*-/-*^ animal and finally DNA from a tumour cell transplanted wild type animal. All scale bars are 50 μm.

### Transplantation validation in irradiated *X. tropicalis* animals

Efficient tumour cell transplantation might also be achieved via alternative techniques apart from the generation of a *rag2*^*-/-*^ line. Immunocompromised *X. laevis* animals can also be obtained by sub-lethal gamma irradiation (Rollins-Smith & Robert, 2019). In order to generate such hosts in *X. tropicalis*, an optimal dose suitable for successful allografting of tumour cells needed to be determined. We irradiated 3 different groups of 4-month-old froglets [8 Gy (n=3), 10 Gy (n=3) and 12 Gy (n=3)] and compared these with a non-irradiated wild type group (n=6) (Fig. 3A). Approximately one week post irradiation all cohorts were euthanized and dissected. Major lymphoid organs (spleen and liver) and peripheral blood were checked to address irradiation potential. Natt and Herrick peripheral whole blood staining revealed significant reduction in white blood cell (WBC)/red blood cell (RBC) ratios in irradiated animals as compared to the non-irradiated controls (*p* = 0.0012) (Fig. 3B). Of note, no significant differences were present between the 3 irradiated groups. Furthermore, quantification of CD3 immunohistochemical stainings revealed that irradiation majorly impacted T-cell levels in both spleens and livers (Fig. 3C-D). For spleens, compared to the wild types (51.9% ± 4.5), irradiation with an 8 Gy dose already induced a significant decrease in CD3 positivity (36.0% ± 5.9, *p* < 0.05). This effect became more pronounced when irradiating to 10 Gy (15.7% ± 2.1, *p* < 0.001) and to 12 Gy (4.9% ± 1.9, *p* < 0.0001). Additionally, in the livers a similar dose-ratio trend was observed [wild type (4.0% ± 1.8), 8 Gy (1.5% ± 0.5, *p* = 0.08), 10 Gy (0.4% ± 0.1, *p* < 0.05) and 12 Gy (0.2% ± 0.1, *p* < 0.05)]. We propose irradiation up to a dose of 12 Gy is preferred for optimal reduction of T-cell numbers, thereby displaying the highest potential for successful tumour transplantation applications.

**Figure 3.**
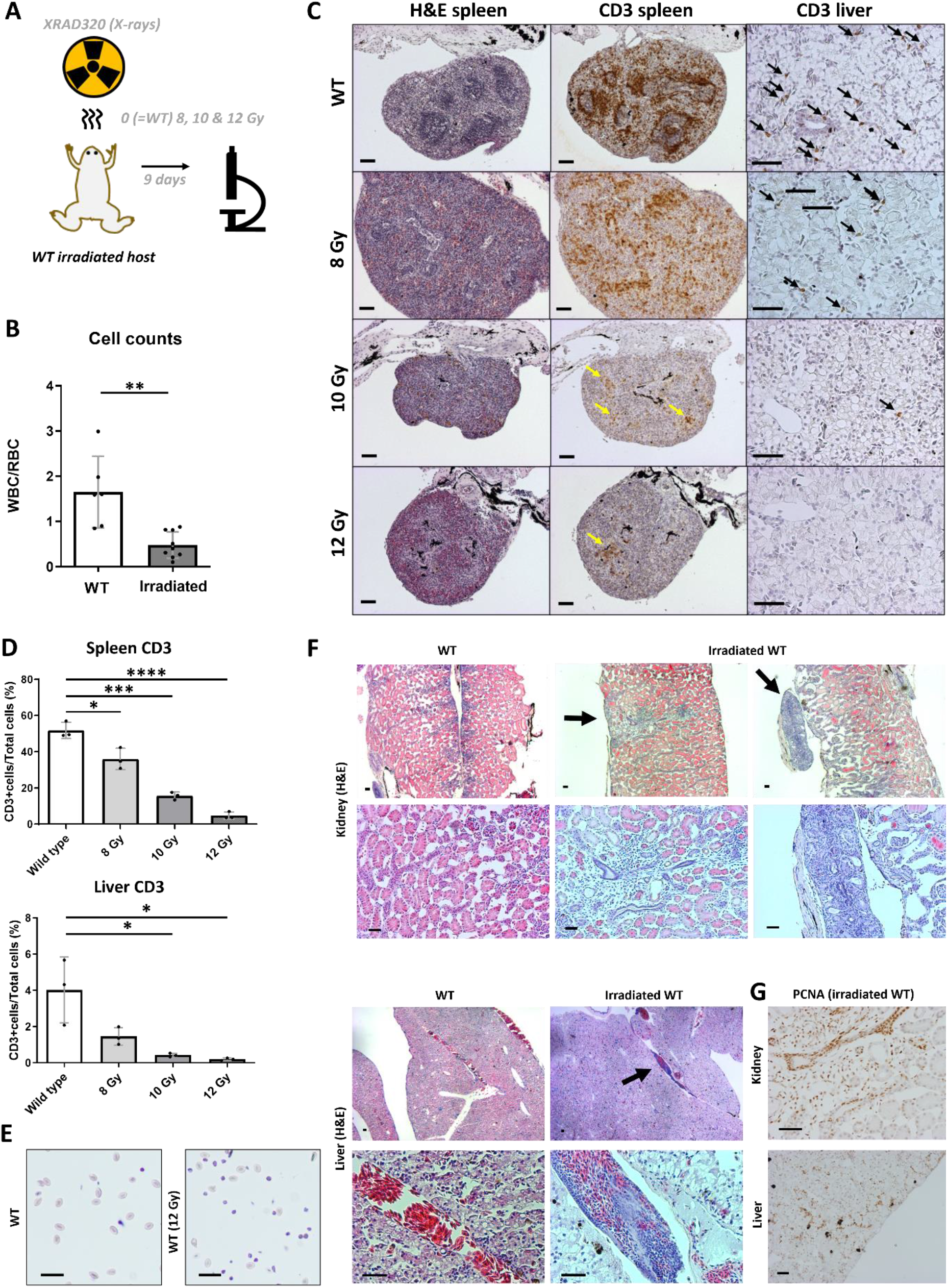
Allografting in irradiated wild type *X. tropicalis* animals. **(A)** Representation of the irradiation procedure for which 3 groups (each n=3) were irradiated with X-rays (8, 10 and 12 Gy) and compared to a non-irradiated wild type group (n=6). **(B)** Plots showing hemocytometer cell counts as represented by white blood cell (WBC)/red blood cell (RBC) ratios of irradiated animals and non-irradiated controls. **(C)** H&E and anti-CD3 immunostained sections from spleens and livers of all 4 groups. Yellow arrows show CD3 positive zones in the spleen, black arrows show CD3 positive cells in the liver. **(D)** IHC quantified CD3 data of spleens and livers using the open source digital analysis tool QuPath (Bankhead et al., 2017). **(E)** IP fluid from transplanted irradiated animal and non-irradiated control stained with Natt and Herrick reagent. **(F)** H&E Sections of engrafted regions in kidney and liver from transplanted irradiated froglet (black arrows) compared to respective kidney and liver sections in the transplanted non-irradiated control froglet. **(G)** PCNA-stained sections from irradiated transplant showing kidney and liver engraftment sites. All scale bars are 50 μm. Bar charts shown represent means with SD as error bar.

In parallel with the experiment in *rag2*^*-/-*^ animals, we validated the transplantation potential of *tp53*-mutant GEXM tumour cells also in an irradiated animal. For this purpose, an irradiated froglet (12 Gy) and a wild type sibling were injected intraperitoneally with 1×10^7^ live tumour cells. To avoid any risk of repopulation of functional immune cells after the irradiation procedure, the froglets were analysed already 3 weeks post transplantation, in absence of any external signs indicative for engraftment. A clear increase of tumour cells circulating in the peritoneal cavity was observed in the irradiated transplant (non-RBC/RBC = 66.7% ± 5.7) as compared to the non-irradiated transplanted control (non-RBC/RBC = 13.9% ± 2.3), where the non-RBC fraction (in the irradiated transplant) was majorly represented by tumour blast cells (Fig. 3E). Furthermore, in-depth histological analysis revealed tumour engraftment in both the kidney and the liver of the irradiated transplanted animal, whereas the wild type transplanted control did not show any signs of engraftment (Fig. 3F). Similar to what was found for the *rag2*^*-/-*^ animal, also tumour grafts observed in the irradiated transplant showed high proliferative capacity as indicated by PCNA immunostaining (Fig. 3G).

## Discussion

Donor cell rejection by the host organism after (allo)transplantation is a common hurdle, jeopardizing *bona fide* assessment of engraftment potential of tumour cells. In absence of syngeneic models, the availability of immunocompromised animals is an absolute need to show evidence of engraftment after transplantation and to allow further phenotypic analysis of cancerous cells.

We describe the generation of the novel *X. tropicalis rag2*^*-/-*^ line as a beneficial tool for transplantation experiments. Due to the central role of the Rag2 protein in the process of V(D)J recombination, these animals should lack mature T- and B-cells. Similar to what has been shown in zebrafish (Tang et al., 2014) also the *X. tropicalis rag2*^*-/-*^ animal used in this study allowed allografting of primary tumour donor cells injected intraperitoneally. Especially for longer incubations and serial tumour transplantations this line is recommended over irradiated animals where the transient nature of the immunosuppression might eventually hamper stable engraftment. Already 10 weeks post transplantation solid tumour grafts were visible at the injection site in the *rag2*^*-/-*^ animal, whereas no signs of engraftment were observed in the control animal. Of note, previous transplantation studies with lymphosarcoma cells in *X. laevis* have shown how infectious Mycobacteria induced granulomas were mistakenly interpreted as the engrafted tumour cells (Asfari, 1988; Asfari & Thiébaud, 1988). Therefore, we would like to state that validation of engraftment should not be solely based on histological assessment. In this study, for example, assessment of engraftment was done via endpoint histopathological analysis with an additional genotypic validation.

Next to mutant or genetically modified hosts, the use of irradiated zebrafish (White et al., 2008) and mouse (Milas et al., 1987) animals have assisted greatly in the cancer research field. For *Xenopus tropicalis* limited data is available that show the potential of using this technique prior to performing allotransplantations. We showed that irradiating froglets with a dose of 12 Gy, reduced T-cell numbers approximately 10-fold in the spleen and 20-fold in the liver. We furthermore showed this dose allowed efficient engraftment of *tp53*^*-/-*^ tumour cells 3 weeks post intraperitoneal injection. Of note, using lower doses of radiation might also be sufficient to allow engraftment of host tumour cells. Goyos and colleagues (2011) showed that a 10 Gy irradiation dose already induced an inhibitory effect on thymocyte survival in *X. tropicalis*.

We hypothesize that engraftment success depends on multiple parameters such as tumour type, number of cells injected, injection site and incubation time in the host. Regarding the latter, it is known that repopulation of functional immune cells in irradiated animals can impair stable engraftment of tumour cells. For example, in zebrafish repopulation of myeloid, lymphoid an immune precursor cells is observed already 2 weeks after irradiating adult zebrafish with 12 Gy (Traver et al., 2004). In agreement with this finding, in another transplantation experiment with *X. tropicalis* GEXM tumour cells in 12 Gy irradiated hosts we indeed observed tumour cell clearance 5 weeks post transplantation, probably due to host immune cell repopulation (manuscript in preparation). Considering this caveat, the availability of the *rag2*^*-/-*^ line offers more flexibility with higher engraftment success rates even for long term experiments. We are convinced that with the generation of our novel *rag2*^*-/-*^ line - and the ease with which irradiation can be performed - studies on immune surveillance and tumour immunity will be significantly aided.

## Material and Methods

### CRISPR/Cas9 mediated generation of mosaic mutant *X. tropicalis* animals

The CRISPRScan software package (Moreno-Mateos et al., 2015) was used for the design of the *rag2* CRISPR sgRNA. A 5’-gaattaatacgactcactataggGTCTTCCCTCCATGAATGgttttagagctagaaatagc-3’ oligo along with the reverse oligo: 5′-aaaagcaccgactcggtgccactttttcaagttgataacggactagccttattttaacttgctatttctagctctaaaac-3′ were ordered (Integrated DNA Technologies). At first DNA was prepared by annealing of the two primers and PCR amplification. The DNA template was *in vitro* transcribed using the HiScribe™ T7 High Yield RNA Synthesis Kit (New England Biolabs). The sgRNA was subsequently isolated using the phenol-chloroform extraction/NH_4_OAc precipitation method (Nakayama et al., 2014). RNA quantity was calculated by Qubit^®^ 2.0 Fluorometer (Thermo Fisher Scientific) measurement and quality was visually confirmed by agarose gel electrophoresis. A detailed guideline for generating the NLS-Cas9-NLS protein can be found in previous described work (Naert et al., 2016). After setting up natural matings resulting 2-cell stage embryos were injected unilaterally with a 1 nl pre-incubated (30 sec @ 37°C) mix of sgRNA and Cas9 protein. Gene editing efficiencies were evaluated quantitatively by targeted amplicon next-generation sequencing (as described below). The inDelphi *in silico* prediction algorithm was included to validate endogenously observed frequencies of variant calls (Shen et al., 2018).

### DNA extraction and sequencing

Gene editing was assessed by subjecting PCR amplified sgRNA targeted regions to deep sequencing followed by BATCH-GE analysis (Boel et al., 2016). DNA, from either whole embryos (three embryo pools each containing three stage NF 41 embryos) or from dissected tumours, was isolated using DNA lysis buffer (50 mM Tris pH 8.8, 1 mM EDTA, 0.5% Tween-20, 200 μg/mL proteinase K) during an overnight incubation (55°C) followed by a 5 min boiling step. Primers used in this study for amplification were: *rag2*^*fw*^ 5’-GCTATCTGCCTCCACTTAGAC-3’ and *rag2*^*rv*^ 5’-AATGTCAATGGTGTCATCATC-3’ with an extra internal primer used for Sanger sequencing *rag2*^*int*^ 5’-TCTCCTATTGACTGAAGATGCC-3’, *tp53*^*fw*^ 5’-CAGTGCTTATTGTTACCTCCA-3’ and *tp53*^*rv*^ 5’-CATGGGAACTGTAGTCTATCAC-3’. The methodology for Sanger sequencing and correlation analysis between *in vivo* versus *in vitro* CRISPR mutational repair outcome can be found in (Naert, Tulkens, et al., 2020).

### (Mixed) HMA genotyping method

For genotyping the *rag2* line and tumour (graft) cells, WT DNA (*i*.*e*. DNA from non-injected frogs) was amplified in parallel with each unknown DNA sample via a standard PCR. Subsequently, equal quantities of both products – PCR amplified WT and unknown sample DNA – were mixed and eventually subjected to HMA in parallel with all the unknown samples individually (unmixed). This was completed by incubation of the samples at 98°C for 5 minutes, followed by a 4°C holding temperature using a transition with a ramp rate of 1°C/s. Finally, the PCR amplicons were prepped with DNA loading dye and run on an 8% (bis)acrylamide/TBE gel. Visualization was done on a Molecular Imager^®^ Gel DocTM XR+ System (Bio-Rad) supported by the Image Lab software (Bio-Rad).

### Irradiation procedure

24 hours prior transplantation, animals (early froglet stage) were sub-lethally irradiated up to 12 Gy with X-rays using the XRAD320 device (Precision X-Ray, Inc, North Branford, CT) at approximately 120 cGy/min. Froglets were placed individually in 50 mL Falcon tubes filled with 25 mL filter sterilised frog water.

### Tumour cell transplantation

Tumour single cell suspensions were prepared manually by dissecting tumour pieces, subsequently washing them with sterile amphibian phosphate buffered saline (APBS) after which they were poured through a 40 μm cell strainer (Falcon™) using tweezers to mince the tumour and APBS for flushing. An aliquoted 20 μL of single cells was mixed with 180 μL 0.1% tryphane blue solution to count living cells. Subsequently, the tumour cell suspension was centrifuged for 5 min at 240 g (RT) and resuspended with APBS to the appropriate concentration. Recipient host frogs (*rag2*^*-/-*^, irradiated or WT) were sedated using a 2 g/L MS222 (Tricaine methanesulfonate) solution diluted in water and adjusted to pH 7 with sodium bicarbonate. Each recipient host animal was injected intraperitoneally with a 100 μL tumour single cell suspension containing 5×10^6^ live tumour cells for the *rag2*^*-/*-^ and respective adult control recipient and 1×10^7^ live tumour cells for the irradiated and respective control froglet recipient, using BD Micro-Fine Demi 0.3 mL Syringes 0.3 mm (30G) x 8 mm. Post transplantation, injected animals were housed separately and monitored closely for any signs of engraftment or discomfort. For all animal experiments, ethical approval was obtained and guidelines set out by the ethical committee were followed.

### Blood counts

Peripheral blood or intraperitoneal fluid was isolated by cardiac puncture or intraperitoneal (IP) lavage, respectively. For the IP lavage, a small incision was made in the skin of the belly and the abdominal muscle wall after which 100 μl APBS was used for rinsing the IP cavity. Approximately 10 μl IP fluid cells diluted in APBS was collected for further processing. Immediately after collection, cells were diluted 1:50 in Natt and Herrick reagent, a methyl violet based staining solution, for downstream counting analysis (Maxham et al., 2016; Natt & Herrick, 1952). Counts were performed using a Buerker hemocytometer (Marienfeld). For each Natt and Herrick sample at least 2×6 regions were counted (minimum 150 cells per count).

### Imaging, histology and immunohistochemistry

Animals were euthanized by lethal incubation in a Benzocaine solution (500 mg/L) until heart beating stopped. Macroscopic images were taken with a Carl Zeiss StereoLUMAR.V12 stereomicroscope. Dissected organs or tumours were fixed overnight in 4% PFA at 4°C and subsequently dehydrated and paraffinized. Organ slices (5 μm) were generated by microtomy and stained with haematoxylin and eosin using the Varistain™ 24-4 Automatic Slide Stainer (Thermo-Scientific) for classical histological assessment. For immunohistochemistry (IHC) experiments following primary antibodies were used: IgG anti-human CD3 antibody (1:200, clone CD3-12, Bio-Rad) and anti-PCNA antibody (1:1000, PC10, Dako). Following secondary antibodies (all 1:500) were used: Biotinylated Goat Anti-Rat Ig (559286, BD Pharmingen) and Biotinylated Goat Anti-Mouse Ig (E0433, DAKO). DAB was used as chromogenic method of detection and signal was developed using the VECTASTAIN Elite ABC HRP Kit (PK-6100; Vector laboratories) combined with ImmPACT DAB Peroxidase (SK-4105; Vector laboratories). Finally, samples were counterstained with haematoxylin. All IHC experiments included ‘no primary antibody’ controls (data not shown). Imaging of sections was performed by using an Olympus BX51 Discussion Microscope. For quantification of CD3 stained slides, the QuPath software tool (Bankhead et al., 2017) was used. Slides were acquired using the ZEISS Axioscan 7 machine at 20x magnification with a resolution of 0.22 μm/pixel.

### Statistical analysis

Comparisons and conclusions between experimental and wild type groups were statistically supported by two-sided student’s t-tests (non-significant *p* ≥ 0.05, \**p* < 0.05, \*\**p* < 0.01, \*\*\**p* < 0.001, \*\*\*\**p* < 0.0001). Bar charts shown represent means with SD as error bar.

## Contributions

D.T., P.V.V. and K.V. designed the study. D.T., D.D., M.B., T.V.N. and S.D. were involved in the generation and validation of the *rag2*^*-/-*^ line. D.T. and W.T. performed the irradiation procedure. D.T. and K.V. performed all transplantations. D.C. performed histopathological validation of the tissue sections. D.T. & K.V. wrote the manuscript.

## Acknowledgements

D.T. holds a PhD fellowship from the Research Foundation—Flanders (FWO-Vlaanderen). Research in the authors’ laboratory is supported by the Research Foundation – Flanders (FWO-Vlaanderen) (grants G0A1515N and G029413N) and by the Concerted Research Actions from Ghent University (BOF15/GOA/011 and BOF20/GOA/023). Further support was obtained by the Hercules Foundation, Flanders (grant AUGE/11/14), the Desmoid Tumor Research Foundation, the Desmoid Tumour Foundation of Canada and SOS Desmoïde. In addition, the authors would like to thank Marjolein Carron and Annekatrien Boel for processing the next-generation sequencing data via the BATCH-GE software. We would like to acknowledge Amanda Gonçalves and Benjamin Pavie (VIB Bioimaging Core) for generating the QuPath script for positive cell detection. Furthermore, we are thankful to Tim Deceuninck for the good animal care. Finally, we would like to thank Joeri Tulkens for critical proof-reading of the manuscript.

